# A microphysiological system for parallelized morphological and electrophysiological read-out of 3D neuronal cell culture

**DOI:** 10.1101/2023.11.06.565744

**Authors:** Peter D. Jones, Beatriz Molina-Martínez, Anita Niedworok, Paolo Cesare

## Abstract

Three-dimensional in vitro models in microfluidic systems are promising tools for studying cell biology, with complex models using multiple cell types combined with high resolution imaging. Neuronal models demand electrical readout of the activity of networks of single neurons, yet classical planar microelectrode arrays struggle to capture extracellular action potentials when neural soma are suspended distant from the microelectrodes. This study introduces sophisticated microfluidic microelectrode arrays, specifically tailored for electrophysiology of 3D neuronal cultures. Using multilayer photolithography of permanent epoxy photoresists, we developed devices having 12 independent culture modules in a convenient format. Each module has two adjacent compartments for hydrogel-based 3D cell culture, with tunnels allowing projection of neurites between compartments. Microelectrodes integrated in the tunnels record action potentials as they pass between the compartments. Mesh ceilings separate the compartments from overlying wells, allowing for simple cell seeding and later nutrient, gas and waste exchange and application of test substances. Using these devices, we have demonstrated 3D neuronal culture, including electrophysiological recording and live imaging. This microphysiological platform will enable high-throughput investigation of neuronal networks for investigation of neurological disorders, neural pharmacology and basic neuroscience. Further models could include cocultures representing multiple brain regions or innervation models of other organs.

## 1 Introduction

Neurological disorders affect more than 90 million people worldwide (World Health Organization, 2006) and the incidence is expected to rise due to population aging. Despite enormous investments in drug development, attrition rates of neuroactive compounds are greater compared to other drugs (Dowden and Munro, 2019). Current in vitro and animal models employed for drug discovery or neuroscience research do not adequately simulate the pathophysiological condition of the human nervous system and have limited readout (Marx et al., 2016).

Traditional two-dimensional (2D) cultures are convenient in vitro platforms for carrying out a detailed analysis of the neuronal response. In 2D cultures, cells grow adherent on planar substrates. These culture models emulate certain aspects of neurophysiology and enable higher throughput and lower complexity than in vivo models. However, planar adherent cultures do not represent the complexity of the neuronal circuits and the intricate connection between the neurons (Marx, 2018). Monolayers of neurons attached to rigid surfaces are limited to contact with surrounding cells and lack the extracellular matrix. In an effort to generate more predictive in vitro models that better mimic aspects of the physiology of the human brain, organ-on-chip technology combining microfluidics and three-dimensional (3D) culture techniques has emerged in the last decade (Mastrangeli et al., 2019).

A key distinction in 3D cultures is whether the cells are self-organized by bottom-up mechanisms or are engineered with top-down mechanisms. Bottom-up neural systems include brain organoids (Lancaster et al., 2013; Wang et al., 2018; McDonald et al., 2023) and neurospheres (Fan et al., 2016). Top-down systems are usually dissociated neurons or other cells distributed in a scaffold. In either case, the cultures may include various types of cells with varying distribution and may include a scaffold or extracellular matrix. In organoids, the types and distribution of cells depends on guiding natural mechanisms of differentiation and proliferation. In contrast, top-down control can directly include mature cell types at desired densities (Jentsch, 2021). Our work presents a top-down approach in which culture conditions and distribution of cells can be defined for specific applications.

Differences between cells cultured in 2D and 3D include levels of gene expression (Kamei et al., 2016; Papadimitriou et al., 2018; Tekin et al., 2018) and electrical activity (Xu et al., 2009). Neurons in 3D can develop a more physiological morphology (Lancaster et al., 2013; Todd et al., 2013). An extracellular matrix or scaffold can support the formation of distinct regions or barriers as in the brain (Wevers et al., 2016). Moreover, microfluidics enables perfusion to imitate the fluid flow, including nutrients, gases and secreted substances (Koo et al., 2018; Papadimitriou et al., 2018; Wevers et al., 2018).

The extra spatial dimension of 3D cultures makes measuring the properties and activity of cells more challenging. Most work examines morphology and relies on calcium imaging to measure activity, without electrophysiology (Hopkins et al., 2015; Mofazzal Jahromi et al., 2019). Unfortunately, measuring the activity of deeper layers of neuronal organoids and 3D cultures remains a challenge and requires new technological approaches (Keller and Frega, 2019). Recent approaches include integration of microelectrodes in 3D arrays using complex assembly methods (Soscia et al., 2020; Shin et al., 2021) or designing 2D arrays to be embedded within 3D cultures (Le Floch et al., 2022; McDonald et al., 2023).

In fact, a promising approach is to develop new microelectrode arrays (MEA) with integrated microfluidic structures (Neto et al., 2016). Direct fabrication of microfluidics on microelectrode arrays allows alignment with optical precision (∼1 µm), and can be scalable. A challenge is the vertical height of required microfluidic structures, which must be >100 µm for 3D cell culture.

Similar work has traditionally relied on placing microfluidic devices made of polydimethylsiloxane (PDMS) on conventional glass microelectrode arrays, defining interconnected compartments for multiple cell populations (Honegger et al., 2013; Odawara et al., 2013; Fan et al., 2016; Kilic et al., 2016; Adriani et al., 2017; Park et al., 2018; Wang et al., 2018; Izzi, 2021; Mateus et al., 2021).

Alignment between the microfluidic device and the MEA often uses flip-chip methods with accuracy of only ∼10 µm. Moreover, PDMS has well known drawbacks for substance testing due to absorption of hydrophobic compounds (Toepke and Beebe, 2006; van Meer et al., 2017). Using 2D adherent cell culture, such PDMS microfluidic chips with interconnected compartments have been used to study axonal transport (Lu et al., 2012), unidirectional axonal growth (Peyrin et al., 2011; Malishev et al., 2015), propagation of action potentials (Pan et al., 2011; Mateus et al., 2021), axonal regeneration and injury (Taylor et al., 2005), cocultures with neurons and glial cells (Adriani et al., 2017; Park et al., 2018) or innervation using cocultures of peripheral neurons with other cells (Uzel et al., 2016).

Towards models with 3D cultures in a scalable format, we have developed a new microphysiological device that enables 3D culture, electrophysiology using integrated microelectrodes and high-resolution optical methods. Microfluidics were fabricated directly on MEAs by photolithography of permanent epoxy-based photopolymers. A notable advantage of this process is the flexibility to build precise multilayer 3D architectures. The inert surface and transparency of the epoxy polymers make them a suitable choice for microfluidic devices and compatible with microscopy techniques (Abgrall et al., 2007; Ren et al., 2013).

With this device, we demonstrated a two-compartment 3D neuronal model, which required low numbers of cells and low medium volumes. Neurons embedded in hydrogel spontaneously extended neurites, including through the interconnecting tunnels with a cross-section of ca. 5×5 µm² (**Figure 1**). After growth of neurites into these tunnels, transmitted action potentials were extracellularly amplified and readily recorded (Molina-Martinez, 2020; Molina-Martínez et al., 2022). We thereby achieved reliable recording of the 3D cultures’ activity using microelectrodes in neurite-trapping tunnels.

**Figure 1:**
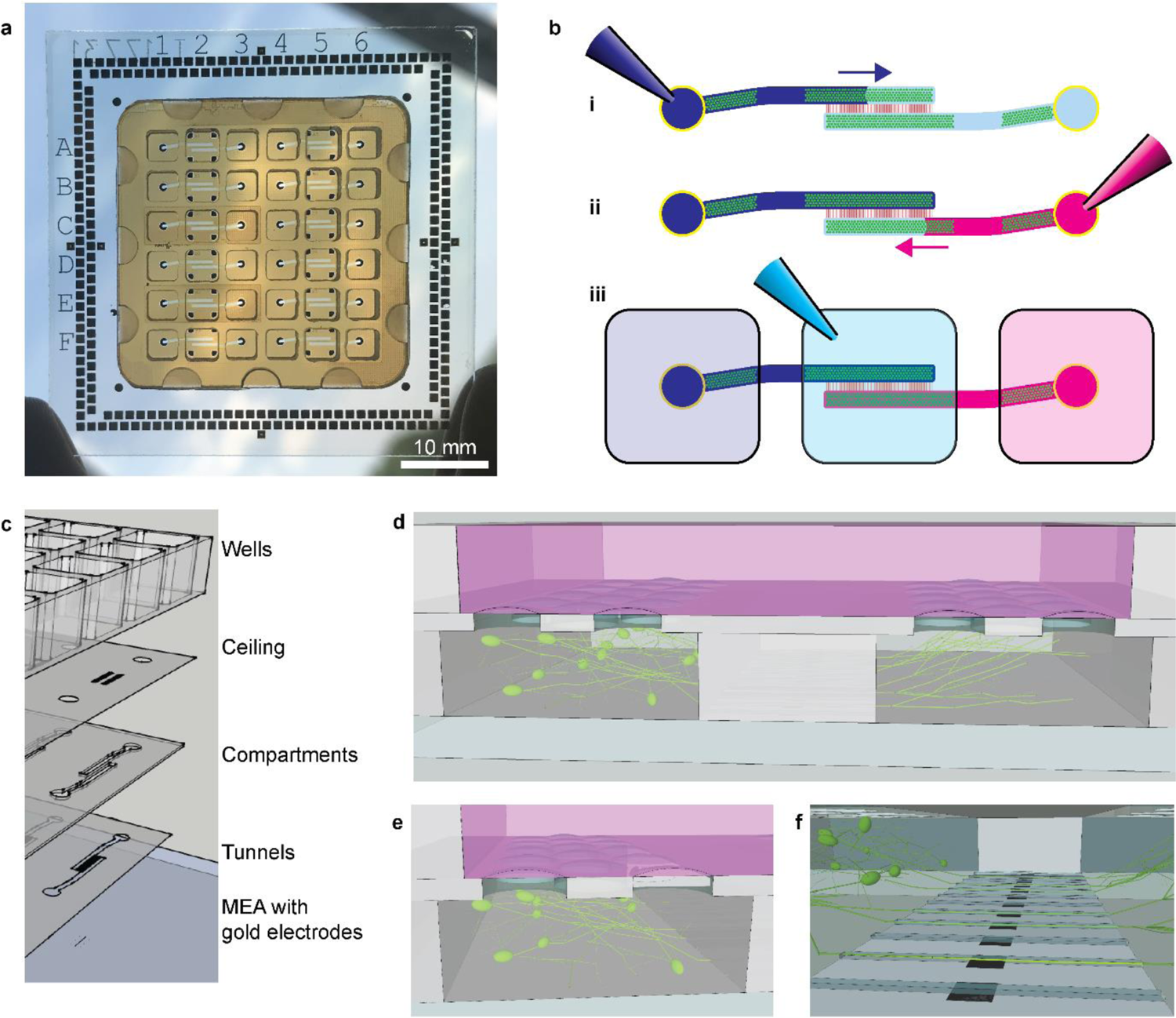
Device concept. **a**: Complete device with 12 microfluidic modules (brown, 32×32 mm²) on a glass-based 256-channel microelectrode array (49×49 mm²). Peripheral contact pads correspond to commercial amplifiers. **b**: An illustration of the filling procedure, with seeding of cells in the two channels (**i**, **ii**) and then filling of the central well (**iii**). Filling with different colors illustrates that different cell populations can be seeded in each compartment. Compartments are connected by tunnels (red) which contain microelectrodes (not shown). The channel ceilings contain a mesh (green) which prevents overflow into the wells during filling. Inlets are outlined in yellow. For clarity, well outlines are shown only in **iii**. **c:** Scheme of the multiple layers for a single module. From bottom to top: a MEA with gold microelectrodes, ITO paths and a silicon nitride insulator; tunnels for neurites (3 µm SU-8); compartments for hydrogel-based cell culture (150 µm SUEX); mesh ceiling for substance exchange (20 µm ADEX); and wells for culture medium (3 mm COC). **d–f:** Cross-section illustrating the two parallel compartments enclosed by a mesh ceiling (d) and connected by the tunnels (e). Neurons are suspended in the hydrogel below the mesh. Media is above the mesh. Neurites project in all directions, including through the tunnels where microelectrodes enable recording of action potentials. **d–f** are snapshots from Supplementary Video 1.

Simultaneously, morphology of the cultures could be observed with high resolution. This method enabled the non-invasive recording of non-adherent neurons with high efficiency and reproducibility, enabling parallelization of up to 12 cultures per device.

## 2 Materials and methods

### 2.1 Microfabrication

The fabrication process is illustrated in Figure 2. Devices were designed with CleWin (WieWeb, the Netherlands). Custom MEAs were ordered from NMI TT GmbH (Reutlingen, Germany). MEAs on glass substrates (170 µm or 1 mm thick) had transparent indium tin oxide (ITO) electrical paths insulated by silicon nitride. After opening microelectrodes by etching the insulator through a photoresist mask, gold was sputter deposited and structured by lift-off. Adhesion of gold directly on ITO was sufficient without an extra adhesion layer.

**Figure 2:**
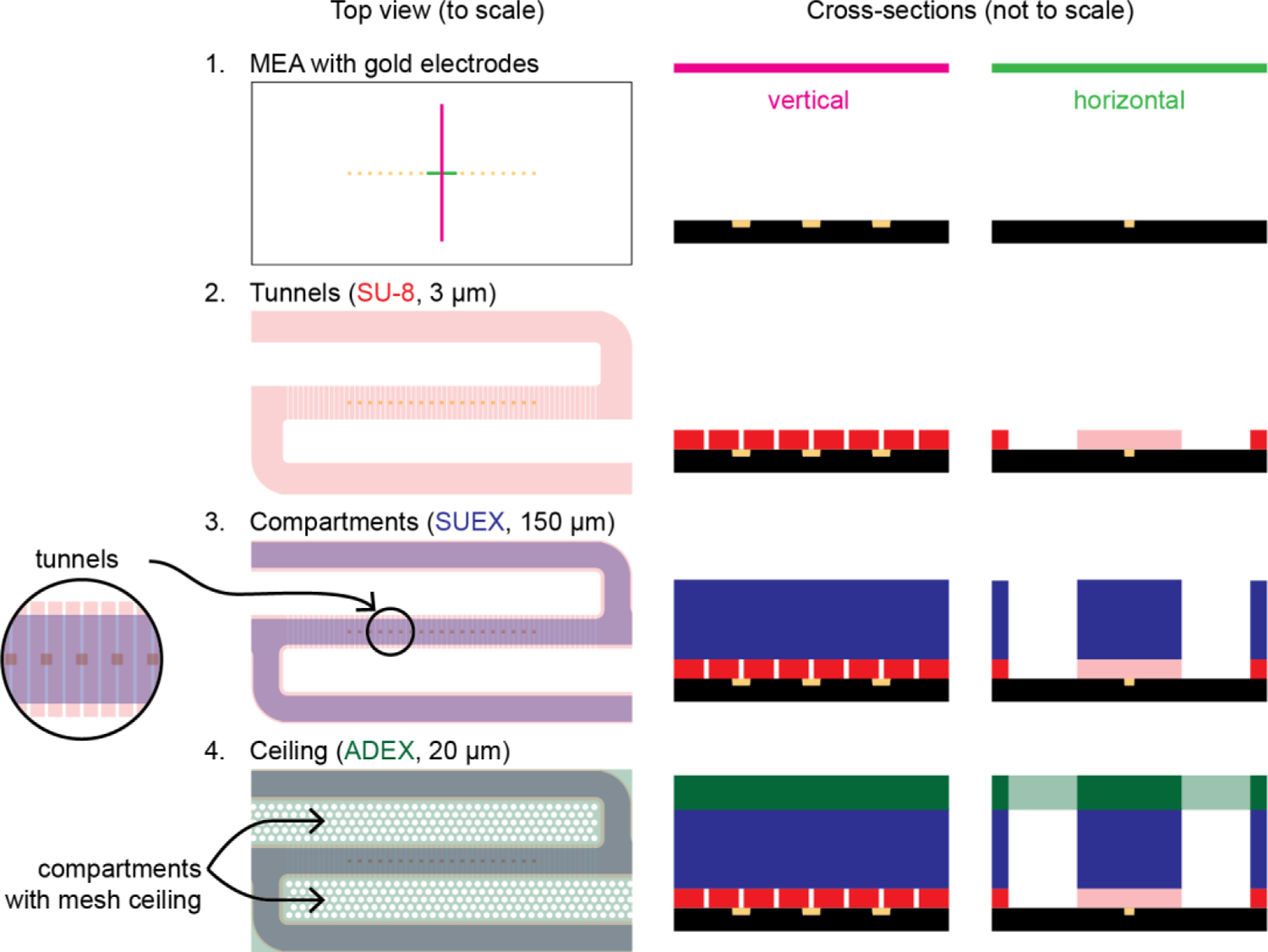
Microfluidic fabrication process. Fabrication begins with a glass substrate with gold microelectrodes (1). The upper surface is silicon nitride, and ITO electrical paths (height ∼200 nm) are omitted for simplicity. Tunnels are produced in a spin-coated layer of SU-8 (3 µm) (2). Compartments are defined by a laminated layer of SUEX (150 µm) (3) and enclosed by a laminated ceiling of ADEX (20 µm) (4).

Microfluidic structures were produced in three photolithographically structured layers of permanent epoxy photoresist, described in detail below and shown in **Figure S1**. The first layer was spin-coated while the second and third layers were dry film resist which was laminated to produce enclosed channels. All baking steps were performed with ramping to and from room temperature, primarily to reduce stress caused by the order-of-magnitude difference between the thermal expansion coefficients of the glass and polymers. We found ramping reduced the need for manual handling as a large hotplate could fit up to 36 MEAs. Heating the hotplate required ca. 10 min, while cooling took more than an hour. For the oven, heating and cooling was controlled at 0.5 °C/min. All UV exposure used an i-line filter. Lamination temperatures were measured by stopping a thermocouple between the heated rollers. Proximity exposure was used to avoid physical damage to dry film resists, which could adhere to the mask and be partially removed.

MEAs were pretreated by oxygen plasma (2 min, 200 W) then baked in an oven for 1 h at 150°C. For the first layer (tunnels), the liquid photoresist SU-8 2002 (Kayaku AM) was spin-coated to produce 3-µm-thick structures (10 s at 500 rpm then 30 s at 1000 rpm). A baking step (5 min at 95°C on a hotplate, ramped) removed solvent. Structures were exposed (250 mJ/cm², ca. 20 s, i-line, soft contact, SÜSS MA6) through a photomask. A post-exposure bake (PEB), identical to the pre-exposure bake, cross-linked the exposed structures. Structures were developed in the SU-8 developer (mr dev 600) for 40 s and then rinsed in isopropanol. An intermediate hardbake (150°C in an oven) further cross-linked the SU-8; this was necessary to avoid overdevelopment of SU-8 in subsequent steps with longer development and cyclohexanone.

For the second layer (compartments), the dry film resist SUEX 150 (150 µm thick) was laminated at 3 mm/s and 78°C. The carrier foil was removed, then SUEX was exposed through a photomask (2500 mJ/cm², i-line, 20 µm proximity exposure). The PEB was 2 h at 45°C, then ramped (ca. 5 min) to 65°C for an additional hour. After cooling, SUEX was developed for 10 min in mr dev 600, rinsed with isopropanol and dried with nitrogen. To remove residues in the tunnels, dry substrates were dipped again in mr dev 600, rinsed with isopropanol and dried with nitrogen.

For the third layer (ceiling), substrates were prepared for lamination by temporarily fixing a 150 µm-thick metal foil spacer around the SUEX structures to create a near-planar surface (**Figure S2**). The dry film resist ADEX A20 (20 µm thick) was laminated at 3 mm/s and 75–78°C. The cover foil was removed and ADEX was exposed (600 mJ/cm², i-line, 20 µm proximity exposure). The PEB was 1 h at 65°C, ramped. ADEX was developed in a two-step process which was necessary to remove unexposed ADEX and clean all residues from the enclosed channels. The first step was dipping in cyclohexanone for 3 min, rinsing with isopropanol for 1 min, and drying with nitrogen gas. The second step was dipping in cyclohexanone for 1 min, rinsing with isopropanol for 10 min, and drying with nitrogen gas. In the first step, cyclohexanone dissolved unexposed ADEX and flowed into the air-filled channels, transporting ADEX residue into the channels; some residue remained after drying. In the second step, fresh cyclohexanone entered the channels and dissolved the residue. In both steps, drying with nitrogen required aiming the nitrogen pistol at the channel openings to clear solvents from them. The metal foil spacer was removed during development.

The microfluidic MEAs were finally hard-baked in an oven at 200°C for 30 min (ramping up and down at 0.5 °C/min) to complete cross-linking for chemical stability (Jones, 2017) and to prevent toxicity (Baëtens et al., 2020).

An array of wells was milled out of COC (Topas 6013) then cleaned in acetone and isopropanol. The microfluidic MEA and wells were treated with air plasma for 90 s, then glued together (**Figure S3**) with EPO-TEK 301-2FL (Epoxy Technology, Billerica, MA, USA) by wicking of the liquid adhesive between the ADEX and COC surfaces (Lu et al., 2008), and cured in an oven at 80 °C for 3 h with ramping up and down at 0.5 °C/min.

### 2.2 Cell culture

#### Preparation of primary mouse hippocampal neurons

Primary cultures were prepared with dissociated hippocampal neurons of Swiss mice (Janvier Labs, France). All experiments were conducted in accordance with the European Union (EU) and German legislation for the care and use of laboratory animals (Directive 2010/63/EU, of the European Parliament on the protection of animals used for scientific purposes, German Tierschutzgesetz).

Isolation of hippocampi from day 16 embryos (E16) was performed based on previously published work (Seibenhener and Wooten, 2012). The cell dissociation procedure followed the guidelines of the Neural Dissociation Kit (T) (130-093-231, Miltenyi Biotec GmbH, Germany). After filtering with medium with 10 % (v/v) fetal bovine serum (10082139, Thermo Fisher Scientific, MA, US), cells were counted (NucleoCounter® NC-200TM, ChemoMetec A/S, Denmark), centrifuged, and resuspended in plating medium based on Neurobasal Plus Medium (A3582901, Gibco, Thermo Fisher Scientific, MA, US) with 2 % (v/v) of SM1 Neuronal Supplement (5711, STEMCELL Technologies Inc., Germany) and 0,5 % (v/v) of Penicillin-Streptomycin solution (P4333-20mL, Sigma-Aldrich, Merck KGaA, Germany).

#### Plating in 3D

Devices were plasma treated (Plasma cleaner/sterilizer PDC-32G, Harrick Plasma, Ithaca, US) for 2 min to sterilize and generate hydrophilic surfaces. As cells should remain suspended in 3D, no coating for cell adhesion was applied. The cell suspension was mixed with a Matrigel matrix (356230, Grow Factor Reduced Matrigel® matrix, Corning Incorporated, MA, US) at a 1:4 ratio in cold conditions and pipetted in the reservoir of each microfluidic channel. The liquid mixture readily fills the channels and compartments, stopping at the mesh ceilings (Figure 1b). Immediately after, the device was incubated upside-down at 37 °C for 10 min to promote the polymerization of the hydrogel and the homogeneous distribution of the neurons. Then, wells were filled with plating medium (50 µL per well). The cell density employed was 10 to 15 thousand cells per microliter and the volume of each channel was 0.4 µl. Cultures were maintained for up to two weeks with medium changes every two to three days. After 6 DIV, plating medium was replaced with maturation medium, consisting of BrainPhys Neuronal (05790, STEMCELL Technologies Inc., Germany) with the same supplements as in the plating medium.

### 2.3 Morphological analysis

One day after plating, neurons growing in 3D were transduced to express a fluorescence protein by adeno-associated virus (AAV) with 10^4^ to 10^5^ virus copies per cell (pAAV-hSyn-hChR2(H134R)-EYFP, 26973, Addgene, MA, US). The expression was monitored by confocal/spinning disk microscopy. Images were taken after one week in vitro with a Zeiss Cell Observer® System with spinning disk head and the software ZEN version 2.3. Afterward, images were processed and reconstructed by Imaris 4.6 (Bitplane AG, Oxford Instruments, UK).

### 2.4 Electrophysiological recording

Spontaneous activity was recorded with a 50 kHz bandwidth using the USB-MEA256 amplifier and MC_Rack version 4.6.2 (Multi Channel Systems MCS GmbH, Germany). Experiments were performed at 36.5 °C, 80 % humidity and 5 % CO_2_, controlled using a customized incubation chamber (Okolab Srl, Italy). Recordings started 10 min after closing the chamber for stabilization of the conditions. Acute response to 3 µM picrotoxin (PTX) was monitored by adding PTX diluted in maturation medium.

Spike detection, burst, and network burst analysis were performed with NeuroExplorer (version 5.212). First, recordings were filtered with a fourth-order bandpass filter (5–6000 Hz). Then, action potentials were detected by a threshold crossing algorithm using five times the standard deviation of the background noise (Quiroga et al., 2004). Burst detection was performed by NeuroExplorer’s Poisson surprise algorithm with a minimum burst duration of 5 ms, a minimum of 5 spikes per burst, and a surprise value of 4. For detection of network bursts, bursts detected on all electrodes (using their median time) were projected as single events in one timeline, and the Poisson surprise algorithm was used with a surprise value of 3 and a minimum duration of 5 ms. Moreover, network bursts were confirmed if at least one third of electrodes (i.e. six) contributed.

### 2.5 Cleaning and reuse

After conducting the experiments, the neuronal culture was discarded and devices were cleaned by rinsing with distilled water and Tergazyme (0.1% w/v) immersion for at least 3 h. After rinsing again with water, drying at room temperature, and plasma cleaning, devices could be reused.

## 3 Results and discussion

### 3.1 Device design

Devices were designed to support simultaneous morphological and functional assessment of 3D neuronal networks. Materials were selected for sufficient thermal stability, solvent compatibility, biocompatibility, and reusability. For the microfluidic structures, permanent epoxy photoresists enabled high resolution patterning of multiple structural layers.

#### Functional structures

Microfluidic microelectrode arrays were designed based on a 49×49 mm² format with 256 electrodes for compatibility with commercial electrophysiology amplifiers (Figure 1a & Figure 4). Microfluidics were addressed by a 6×6 array of wells with a 4.5 mm pitch based on the ANSI/SBS 384-well microplate format for compatibility with automated liquid handling.

Each device includes 12 separate modules for independent experiments. A single module has a 3-well footprint of 4.5×13.5 mm². Two channels (width 400 µm and height 150 µm) with inlets in the outer two wells meet in the central well (Figure 3f**/g**). Each channel defines a compartment for 3D cell culture.

**Figure 3:**
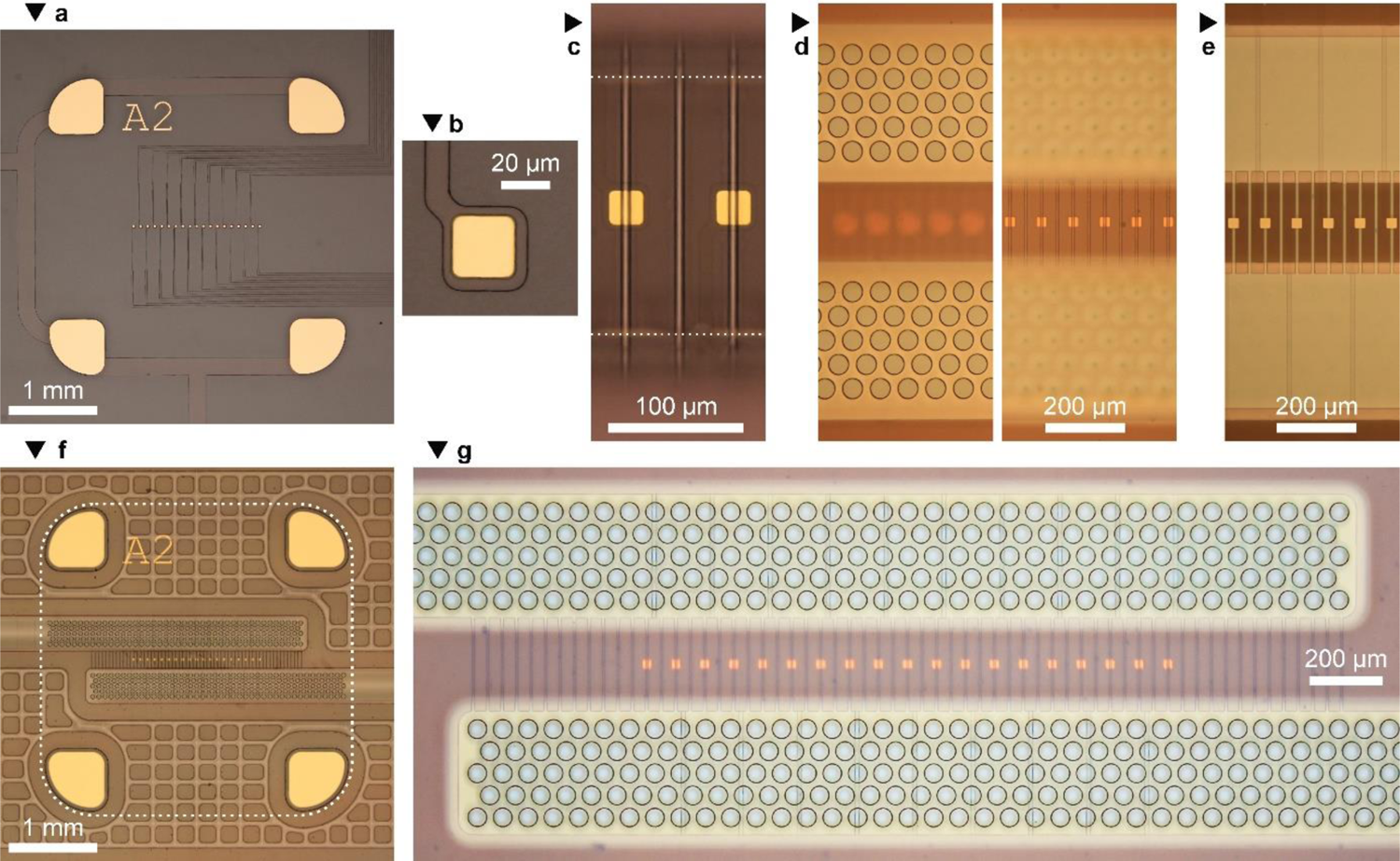
Microfluidic features. **a**: The 19 microelectrodes and segmented reference electrode of a single module. Electrodes are addressed by transparent insulated paths on a glass substrate. **b:** A single microelectrode is 25×25 µm². **c:** Tunnels are defined by the SU-8 layer (3 µm) and the SUEX layer (150 µm). The tunnels are 7 µm wide. The tunnel length of 200 µm is defined by SUEX. The vertical SUEX walls create visible reflections of the tunnels (above and below the added dotted white lines). **d:** The channels are covered by a mesh (pore size 50 µm in a 20-µm-thick ADEX layer). Different focal planes resolve the mesh (left) and the tunnels (right). **e:** Observing the device from below allows the tunnels, electrodes and electrical paths to be resolved at high resolution. SU-8 structures extended 30 µm from both tunnel ends. **f:** The central well of a single module. Segmented structures surround the channels to avoid stress-induced delamination. Not shown is the well (indicated by a dotted line). **g:** The central region of each module comprises two opposing channels connected by 61 tunnels containing 19 microelectrodes. Three images at different focal planes are merged to show all features. Except for **e**, all images were taken from above.

**Figure 4:**
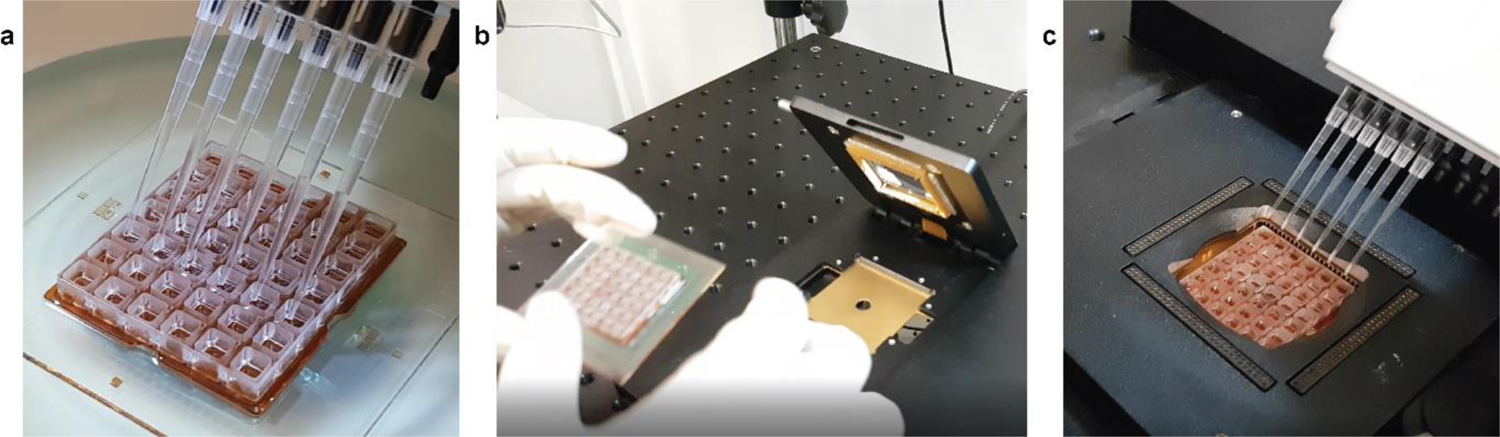
Handling of the microfluidic device. **a**: The 36 wells were designed for handling with multi-pipetting systems for cell plating and medium exchange. **b**: The layout was compatible with 256-channel amplifiers from Multi Channel Systems. **c**: Application of compounds to the 3D neuronal circuits while recording functional effects.

Cells were seeded by pipetting a liquid hydrogel mixture into each inlet, filling each respective compartment and stopping at the mesh (Figure 1b). The two compartments are interconnected by 61 perpendicular tunnels (width 7 µm, height 3 µm, length 200 µm, pitch 40 µm). The ceiling of each compartment is perforated with 50 µm pores (Figure 3d) which constrains flow while facilitating substance exchange (no active perfusion was necessary). This design allows separate seeding of cells suspended in hydrogel in each compartment. The mesh ceiling acts as a capillary stop during seeding, allowing a consistent volume of hydrogel with suspended cells to be seeded without overflowing the compartment. Their culture is restricted so that cell bodies remain within each compartment, while the outgrowth of neurites through tunnels to form inter-compartment synaptic connections is possible.

Each module includes 19 microelectrodes located individually within 19 of the 61 tunnels. Each microelectrode has a square gold surface of 25×25 µm² (Figure 3b), although only a 7×25 µm² region is exposed within the tunnel (Figure 3c). These microelectrodes record action potentials travelling through entrapped neurites (Figure 1d-f). The restriction of a tunnel amplifies the extracellular potential generated around neurites to measurable levels (Molina-Martínez et al., 2022). We note that action potentials were easily detected despite using gold electrodes, which are typically inferior due to their high impedance (Gerwig et al., 2012), and additional noise added by the tunnels’ resistance (Mierzejewski et al., 2020). Gold was chosen due to its electrochemical stability and resistance to contamination. Electrode materials with lower impedance would be TiN or PEDOT:PSS (Mierzejewski et al., 2020), but these would complicate the fabrication. The central well contains a reference electrode (segmented in the well’s corners). The outer wells include electrodes for impedance measurements, which have not been used in the current work.

#### Ancillary features

Large structural areas of thick epoxy resist were segmented to reduce stress and prevent delamination at the polymer/glass interface. Specifically in the 150-µm-thick SUEX layer, features had a maximum lateral size of 200 µm and were separated by 50 µm gaps, creating a pattern of walls and pillars aligned to a 250 µm square grid (Figure 2). The gap width limited sagging of ADEX during lamination and baking steps, while being easily resolved during the UV exposure (aspect ratio height/width of 3). Without these stress-release features, large areas of thick resist were successfully structured but delaminated from glass upon cooling from the final hard bake (**Figure S6**).

While solving the stress problem, this solution created an air-filled cavity within the device. Any defects in the SUEX walls or ADEX ceiling allowed infiltration of the developer cyclohexanone into the cavity. Trapped cyclohexanone swelled and deformed the epoxy structures. To mitigate this problem, walls in the SUEX layer compartmentalized the cavity for each module of the device, so that infiltrated solvents would only affect single modules. An improved design would keep the stress-release cavity open, so that solvents could be easily rinsed. The open stress-release cavities could be filled at a final stage, for example by a potting silicone.

### 3.2 Microfabrication

Glass substrates with gold electrodes, ITO electrical paths, and a silicon nitride insulator were produced by NMI TT GmbH. Each substrate contained twelve modules (Figure 1a), each of which had a row of 19 microelectrodes and a segmented reference electrode with a total area of 1.3 mm² (Figure 3a).

Microfluidics were produced on the glass substrates by photolithographic structuring of epoxy photoresists (Figure 1c). Successful fabrication required optimization of pretreatment, UV dose, baking and developing. Resolution depended on appropriate UV doses. Over-exposure increased structure sizes, which was critical if tunnels became blocked. Adhesion at the silicon nitride/SU-8 interface was excellent with pretreatment by oxygen plasma and baking, which can be understood by cross-linking between a (3-glycidyloxypropyl)trimethoxysilane in SU-8 with silanol groups on the silicon nitride surface. In contrast, adhesion between photoresist layers is explained by epoxide reactions; each processed layer includes residual epoxide groups until final hard baking. In fact, oxygen plasma treatment between successive epoxy layers led to adhesive failure.

The hard bake was important for adhesive stability of the final microfluidic structures in addition to its importance to eliminate toxicity (Baëtens et al., 2020). Slow ramping was critical to prevent thermal stress. The thermal shock of removing devices from a hot oven caused delamination accompanied by audible cracking. In a rare case, we observed that thermal stress caused cohesive failure within the glass substrate.

Lateral segmentation of the 150-µm-thick SUEX to reduce stress was critical for successful fabrication. Glass and epoxy photoresists have a large difference in coefficients of thermal expansion (<10 and 50–100 ppm/°C, respectively), so that a conservative estimate of the extra expansion of the polymer at 200 °C is 0.7 %. A 32-mm-wide structure would need to accommodate 230 µm of strain. In contrast, the segmented structures had at least one dimension limited to 200 µm, which would need to accommodate a strain of 1.4 µm.

Inspection of thick structures revealed peculiarities not observed in thin films. The vertical walls of SUEX created reflections (Figure 3c). When observed from above, the tunnels and microelectrodes (observed through 150 µm of SUEX) and electrical paths (observed through air-filled channels) were at different focal planes. These issues hindered interpretation of micrometer-scale features at the tunnel openings. Observation from below (Figure 3e) allowed resolving these features in a single focal plane. Filling the channels with isopropanol was a simple way to demonstrate that tunnels were unblocked (as isopropanol could be observed entering and flowing through the tunnels).

In processing dry film resists, we used UV doses of two to three times the recommended values and lower PEB temperatures; these changes prevented collapse of tented structures. For the SUEX layer, the first step (45 °C) of the PEB enabled reflow of un-exposed SUEX into the tunnels while exposed SUEX remained stable. The second PEB step at 65 °C cross-linked exposed SUEX. Thereafter, unexposed SUEX could be developed and tunnels were unblocked. Skipping the first step caused reflow of exposed SUEX into the tunnels and its subsequent cross-linking created permanent blockages. Despite optimized processes, a handful of tunnels (∼2 %) were observed to be blocked in finished devices (**Figure S4**). These blockages may be due to shadow-like defects in SUEX as revealed by scanning microscopy (**Figure S5**).

The ADEX layer (20 µm thick) formed tented structures across channels (400 µm) and stress-release features (50 µm). The higher UV dose and lower PEB temperature reduced sagging of these tented structures. Sagging occurred not only during PEB, but also the hard bake. We observed that ADEX spanning stress-release features bulged upwards by 1 µm due to expansion of enclosed air during baking, while regions above the open-ended channels sagged downwards by several micrometers (**Figure S5c**).

### 3.3 Device usability

Our device was designed for compatibility with multipipetting and automated liquid handling systems (Figure 4a**/c**). This feature ensured faster and more precise medium changes and compound application in the twelve modules. While this MEA has a 256-electrode layout compatible with commercial amplifiers (Figure 4b**/c**), the modular design could be adapted for other amplifier formats such as multiwell systems. In this 12-module design, each module requires three wells of the 36-well layout. In previous work, we have reported on the experimental variability in 12- and 18-module designs (Molina-Martínez et al., 2022). The number of modules, the number of electrodes per module, as well as the total number of electrodes and the connector layout are adaptable. Our 36-well layout has a pitch of 4.5 mm (matching 384-well microplates and enabling use of related hardware).

The transparent glass substrate below the 3D cultures enabled optical microscopy. Furthermore, the photoresists, adhesives and polymer wells proved to be biocompatible. These materials were easily sterilized and hydrophilized after plasma treatment, which supported the culture of neurons embedded in the hydrogel. Lastly, cleaning of the device with a detergent solution removed cells and hydrogel and enabled the reuse of devices up to 10-15 times until defects in microfluidic features were observed.

### 3.4 Growth and morphology of 3D neuronal circuits

With our device, we demonstrated 3D culture of neurons forming functional neuronal networks within and between the two compartments (Figure 5a). The top-down approach enabled control of cell types and densities, in contrast to bottom-up organoids or neurospheres. The 3D cultures with a thickness of 150 µm enclosed by the perforated ceilings were healthy without active perfusion or agitation, and culture medium in the wells could be exchanged without disturbing the cultured cells. This method avoided complicated microfluidic tubing systems, which can have issues of bubbles and contamination. Furthermore, perfusion systems can require larger volumes of culture medium.

**Figure 5:**
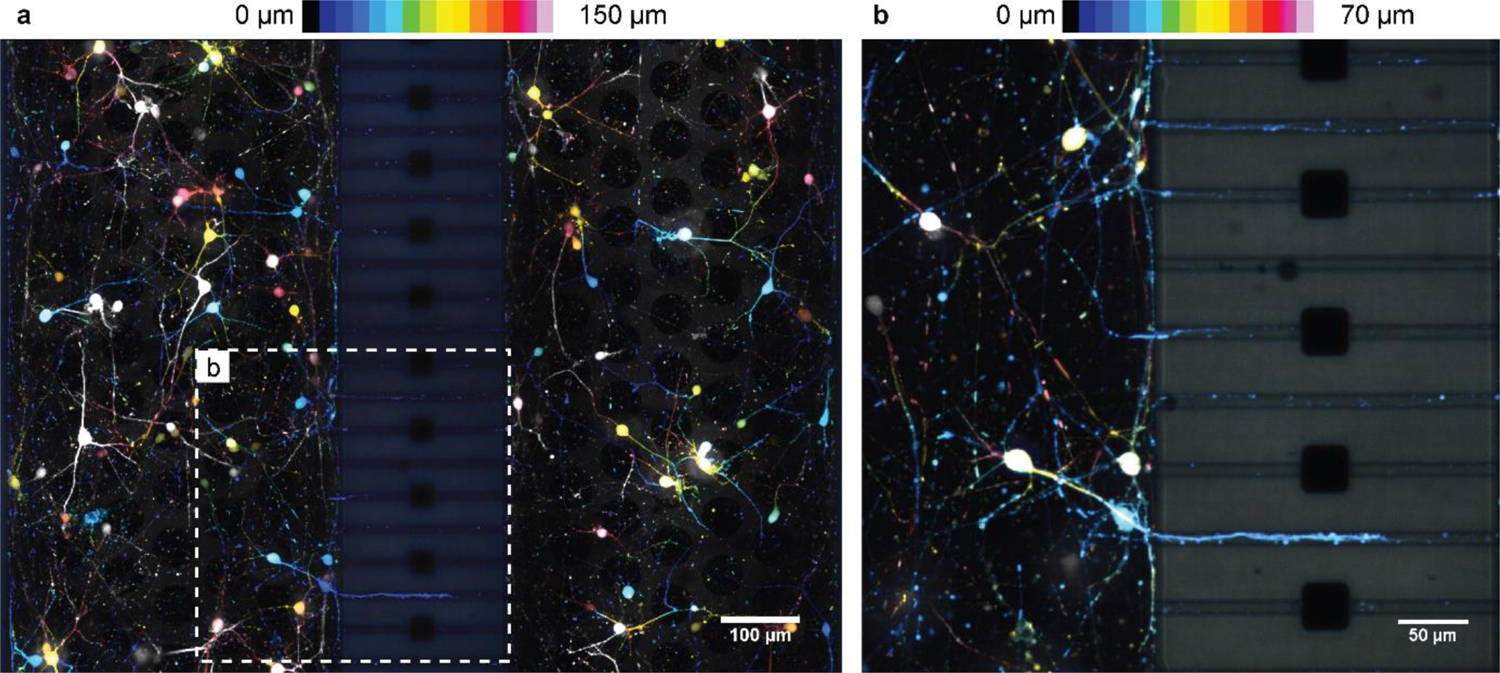
Neurons cultured in 3D extend neurites through tunnels. Neurons (13 DIV) expressing GFP allowed high resolution live imaging of their morphology. Images captured at multiple focal planes within the 150 µm hydrogel thickness are rendered as maximum intensity projections with depth encoded as color. Neuronal soma were suspended in the hydrogel and neurites extended through the tunnels, enabling electrophysiological recording by the integrated microelectrodes. The magnification in **b** focuses on neurites extending into the tunnels. The structure of the tunnels is visible due to autofluorescence of the epoxy photoresist. Gold microelectrodes appear black.

Primary neurons mixed with the hydrogel were distributed in the 3D space and soma remained suspended in the hydrogel without adhering to the device. After four to five days, neurons were live-labeled with GFP and morphology could be monitored by confocal microscopy. Neuronal populations in the two compartments were interconnected by neurites through the tunnels (Figure 5b) ensuring that functional activity would be recorded by the integrated microelectrodes (Figure 6). By employing confocal microscopy, the impact of the autofluorescence of the epoxy structures was minimized, and subcellular features were visible even in the narrow tunnels.

**Figure 6:**
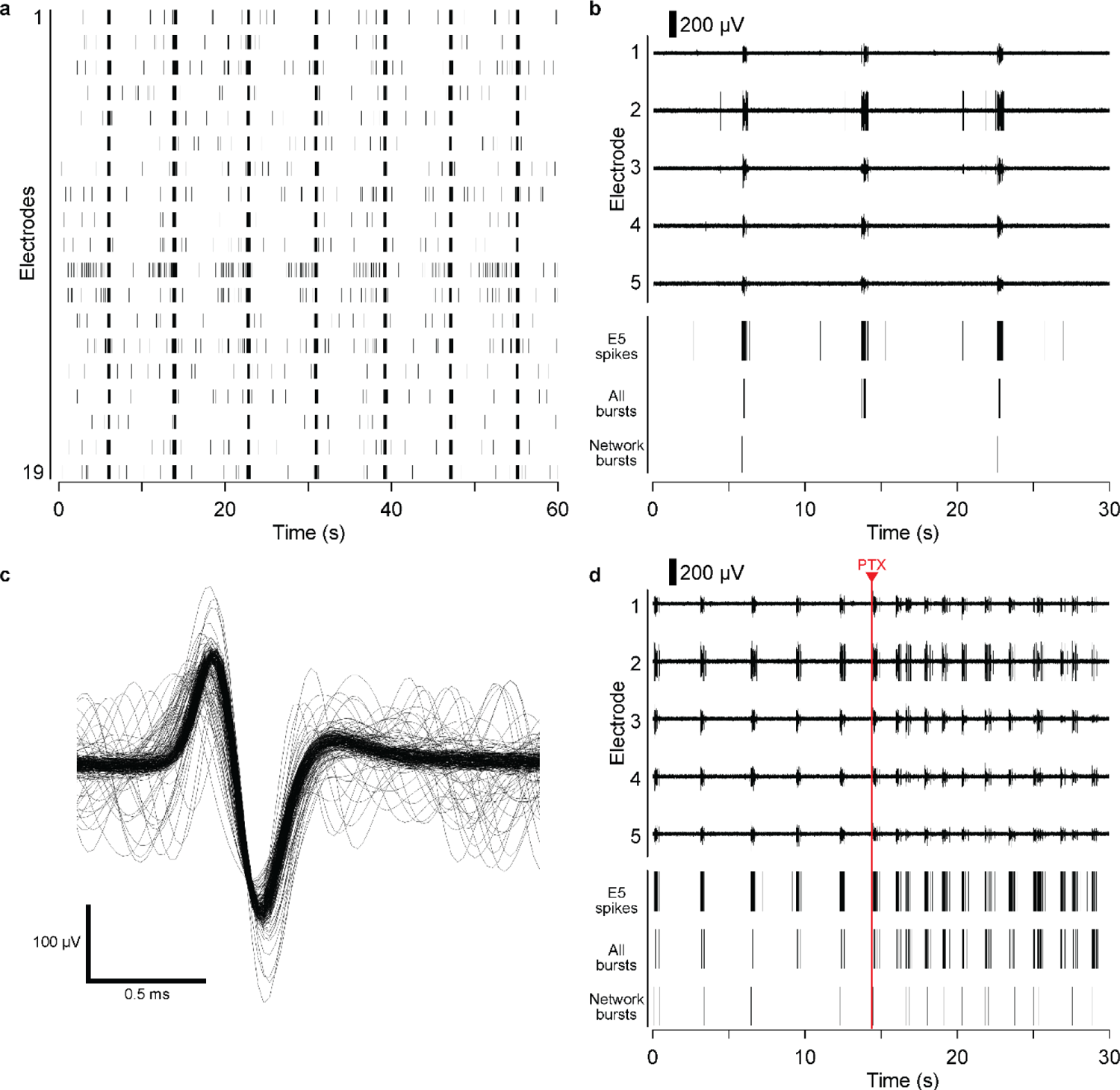
Electrophysiological recordings of 3D neuronal cultures after 13 DIV. **a**: A representative 1 min raster plot of spontaneous activity captured by 19 electrodes in a single module. Each spike is represented by a vertical line. Bursting occurred roughly every 5 s synchronized on all electrodes. **b:** Representative voltage traces of five electrodes, with raster plots of spikes at one electrode, bursts from all electrodes, and network bursts. **c:** Threshold-aligned waveforms of 100 spikes measured at a single electrode revealed the expected waveform of an extracellular axonal action potential, with an amplified amplitude (here, 330 µV peak-to-peak). **d:** Acute application of 3 µM PTX increased spike frequency, illustrated by five representative voltage traces, and raster plots of spikes at one electrode, all bursts, and network bursts.

We observed shrinkage of the hydrogel scaffold over days in culture, resulting in a visible gap forming between the channel walls and the hydrogel (with embedded cells). This shrinkage appeared to be caused by the growing neurons and was proportional to the neuronal cell density; without plating cells, hydrogel did not shrink. With a density of 10 to 15 thousand cells per microliter the 3D culture could be maintained for up to two weeks. Lower densities would inhibit network formation and reduce the likelihood of neurites entering the tunnels.

In our work, the same cells were cultured in both compartments of each module, as our focus was on demonstrating successful cultures with morphological and functional readout. However, the design of this device encourages the combination of two different cultures. This capability could allow the investigation of different neural types, analyze the transfer of protein deposits associated with neurodegenerative disorders between two neuronal populations (Dujardin et al., 2014; Song et al., 2014) or study innervation by including non-neuronal cells in one compartment. Moreover, by addition of endothelial cells directly in the central well of each module, a blood-brain model could be developed. Transendothelial electrical resistance (TEER) measurements would be possible between the reference electrodes in the central well and large electrodes in each outer well (visible at all inlets in Figure 1a).

### 3.5 Electrophysiology of 3D networks

Electrophysiology is essential for analysis of neuronal cultures and for observing the functional effect of compounds, yet the challenge of electrophysiology in 3D cultures presents the need for new technology. We have previously reported a solution using neurite-trapping microelectrodes (Molina-Martínez et al., 2022). This work presents a variation in which the neurites extend between separate culture compartments. In this device, we integrated 19 microelectrodes per culture module, with 12 modules per device having a total of 228 microelectrodes. The recording capabilities rely on the unguided projection of neurites into the tunnels.

Figure 6 illustrates an example of spontaneous activity recorded after 13 DIV. In contrast to patch-clamp (Irons et al., 2008) or conventional “open” microelectrodes (Soscia et al., 2020), the likelihood to record in this system relies on the growth of neurites through the tunnels with integrated electrodes and not on the distance of the soma to the sensor. The chance of an “open” microelectrode (that is, not in a tunnel) of being so close to the soma to record an extracellular action potential is low (Molina-Martínez et al., 2022). In Figure 6a, all 19 microelectrodes in a single module recorded action potentials from 3D cultured neurons. This high capture efficiency was typical in our cultures; neurites crossed most tunnels after one week in vitro despite the outgrowth being spontaneous and unguided.^1^

Recorded spikes demonstrated synchronized behavior, as indicated by bursts recorded simultaneously on all electrodes (Figure 6b). The development of spontaneous activity was comparable to 2D cultures (Biffi et al., 2013; Bradley and Strock, 2019). This synchronized firing behavior demonstrated that neurons have formed a mature and synaptically connected network.

Recapitulating the amplification mechanism of the tunnels (Molina-Martínez et al., 2022): An intracellular action potential is generated by transmembrane ionic currents, and these currents generate extracellular potentials as they disperse through the conductive extracellular space.^2^ When a neurite is trapped in an insulating tunnel, the dispersion of current is restricted (i.e. the current experiences a higher resistance) and thereby the resulting voltage amplitude is higher. This effect amplifies the extracellular action potential of a neurite to levels that can be resolved against the background noise of several microvolts. The recorded waveforms are as expected for extracellular axonal action potentials (Figure 6c).

To demonstrate the capability to monitor the effect of chemical compounds on 3D cultures, we applied picrotoxin (PTX) during recordings. PTX is an antagonist of gamma-aminobutyric acid receptors and a hyperactive reference substance. Acute application of 3 µM PTX rapidly modified the electrical activity inducing more frequent and regular synchronous activity (Figure 6d). This effect corresponds to the expected response of a mature neuronal network response with inhibitory neurons after 13 DIV (Bradley and Strock, 2019). Moreover, these data demonstrating the chemical modulation of 3D neuronal cultures show that PTX diffused through the hydrogel within seconds, as expected for small molecules.

It is important to note that in this device neurites may come from either of the two compartments, and that multiple neurites from multiple neurons may enter single tunnels. Unidirectional outgrowth could be achieved by structured tunnels (Holloway et al., 2019). Different waveforms can be recognized in some cases, although we have not investigated this in detail. If activity is synchronized as in our recordings, the multiple neurites may generate overlapping spikes. Moreover, the integration of two microelectrodes in a single tunnel could determine the direction and speed of action potential propagation.

We have demonstrated a reliable approach to record the activity from 3D neuronal circuits by spontaneous trapping of neurites in tunnels having integrated microelectrodes. Contrary to other techniques, recording does not depend on the location of the neuronal soma in the 3D space. Our approach with substrate-integrated microelectrodes should prove useful for various 3D neural cultures.

## 4 Conclusions

Here, we reported on the development of a two-compartment microphysiological system for the morphological and electrophysiological assessment of 3D neuronal cultures, and demonstrated its capabilities with primary 3D cultures of mouse hippocampal neurons. We developed this system to be compatible with automated liquid handling and commercial amplifier systems, although we note that the culture modules (with 12 in our format) can be extended to multiwell format devices with higher numbers of modules and electrodes.

Our devices reliably achieved simple and non-invasive recording of electrical activity due to the unguided outgrowth of neurites from non-adherent cells into substrate-integrated tunnels having integrated microelectrodes. The devices were also capable of simultaneous microscopy to observe morphology of the 3D neuronal culture.

We demonstrated the formation of mature neural networks within two weeks and modulation of their activity by acute application of a neuroactive compound. The high percentage of active electrodes (up to 19/19) in each module suggest that more modules with fewer electrodes could be integrated per device, which would decrease the cost per data point. Our design without the need for active perfusion presents a simple system for neural organ-on-a-chip models.

With the functionality of the system demonstrated, future work can increase the biological complexity for specific applications. As the system includes two separate culture compartments, two different neuron types could be cultured. Cocultures with other cell types could increase the complexity of the system. One compartment could be used for a non-neuronal culture to study innervation. Similarly, endothelial cells could be cultured above the ceiling in the central well as a model of the blood-brain barrier.

While our devices proved to be reusable, future technical developments could include developing scalable fabrication methods using other materials to reduce fabrication complexity. While we note that our system provides capabilities that are not commercially available, adoption of such devices for commercial purposes will depend on achieving fabrication with a quality and cost that are sustainable for producers as well as customers.

## Supporting information

Supplementary Video 1: Cross-section of the device concept

Supplementary Video 2: Neurite outgrowth through tunnels

## 5 Author contributions

PDJ, BMM and PC conceived and designed experiments. BMM, AN and PC performed and analyzed biological experiments. PDJ developed fabrication methods, designed and produced devices. PDJ and BMM wrote the manuscript. All authors edited the manuscript and approved its final version.

## Acknowledgments

We acknowledge financial support from the State Ministry of Baden-Wuerttemberg for Economic Affairs, Labour and Tourism, and from the German Ministry of Education and Research (BMBF) under Grant Agreement 031L0061 (MEAFLUIT).

## Supplementary Material

Figure S1: Layer structure of the microfluidic module.

Figure S2: Lamination process for dry film resist on non-planar topography.

Figure S3: Gluing of wells to microfluidics.

Figure S4: Yield of tunnel fabrication.

Figure S5: False color scanning electron microscopy images of thick epoxy microfluidic structures.

Figure S6: Delamination of unoptimized epoxy microfluidics from glass.

Figure S7: Neurite outgrowth through tunnels. Supplementary Video 1: Cross-section of the device concept. Supplementary Video 2: Neurite outgrowth through tunnels.

## Footnotes

1 Supplementary Video 2 (**Figure S7**) is a time-lapse video of neurite outgrowth over two days. The microfluidic device has similar dimensions but is made of PDMS on glass. Neurons were cultured in 2D to enable unlabeled imaging.

2 The dispersion of currents and the resulting extracellular potentials are known to depend on the local electrical resistivity, which is higher in tissue than in cell culture medium. Neither enables extracellular recording of axonal action potentials. However, the resistivity of our polymer structures surpasses these by many orders of magnitude.

