## Supplemental Figures S1-S7 for "A microphysiological system for parallelized morphological and electrophysiological read-out of 3D neuronal cell culture"

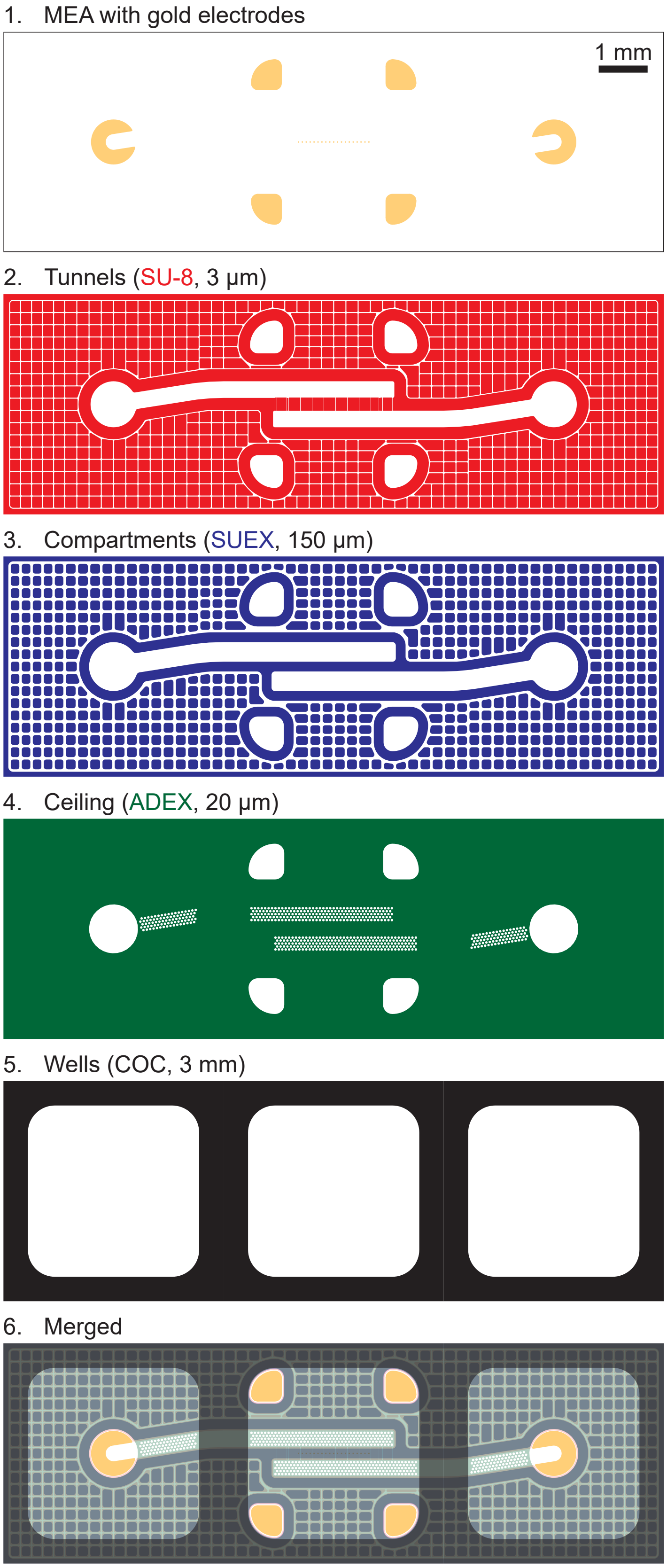


**Figure S1: Layer structure of the microfluidic module.** The devices consisted of a stack of layers, beginning with a MEA having gold microelectrodes (1), followed by three layers of epoxy photoresist (2–4) and topped by milled thermoplastic wells (5). The merged image (6) shows the finished arrangement corresponding to **Figure 3g**.

**Figure S2: Lamination process for dry film resist on non-planar topography.** Laminating onto thick structures with vertical steps is challenging. Here, 3 µm SU-8 and 150 µm SUEX were patterned in a 32×32 mm² region on a 49×49 mm² SiN-coated glass substrate. Simple lamination of the next layer would damage the dry film resist, trap air under the resist and worsen adhesion. **a:** A metal foil spacer with a thickness of 150 µm was laser-cut to match the shape of the microfluidic structure, aligned, and fixed with Kapton tape on two sides. **b:** The surfaces of SUEX and the spacer were nearly coplanar, with a small gap between them. **c:** A piece of 20 µm ADEX with cover foil on both sides was cut to size using a scalpel, aligned, and fixed with Kapton tape on one side. We marked the top cover foil (concave side) with “A20” and the bottom cover foil with “R” so that a quick glance can identify whether a foil has been removed. **d:** ADEX was folded back and the bottom foil was removed with tweezers. By slightly damaging the corner of the ADEX, we visually confirmed removal of the bottom foil while ADEX remained on the top foil. **e:** The laminator temperature was adjusted using an external thermocouple. **f:** A card stock sleeve was stopped between the rollers, and the substrate was placed on the sleeve with the ADEX folded up. **g:** The laminator was started. As the substrate entered the rollers, ADEX was held up gently by tweezers. **h:** After lamination, the substrate was allowed to rest for several minutes. Then the top cover foil was peeled off using tweezers. If the foil was peeled off immediately after lamination, the ADEX was often pulled off the substrate. Peeling back the foil with a large radius of curvature rather than pulling it up sharply was important to prevent pulling off parts of the ADEX from the substrate. **i, j:** Good adhesion of dry film resist was observed by dark contact areas, while suspended regions, poorly adhered regions and trapped air were bright. **k, ℓ:** Substrates were then exposed and underwent a post-exposure bake. During the PEB, exposed regions polymerized and remained adherent to the underlying structures. Unexposed regions, including the suspended ADEX between the microfluidic structure and spacer, remained soft and could deform, as shown here after removal of the Kapton tape relaxed the spacer. **m:** During immersion development, spacers fell off as the unexposed ADEX dissolved.

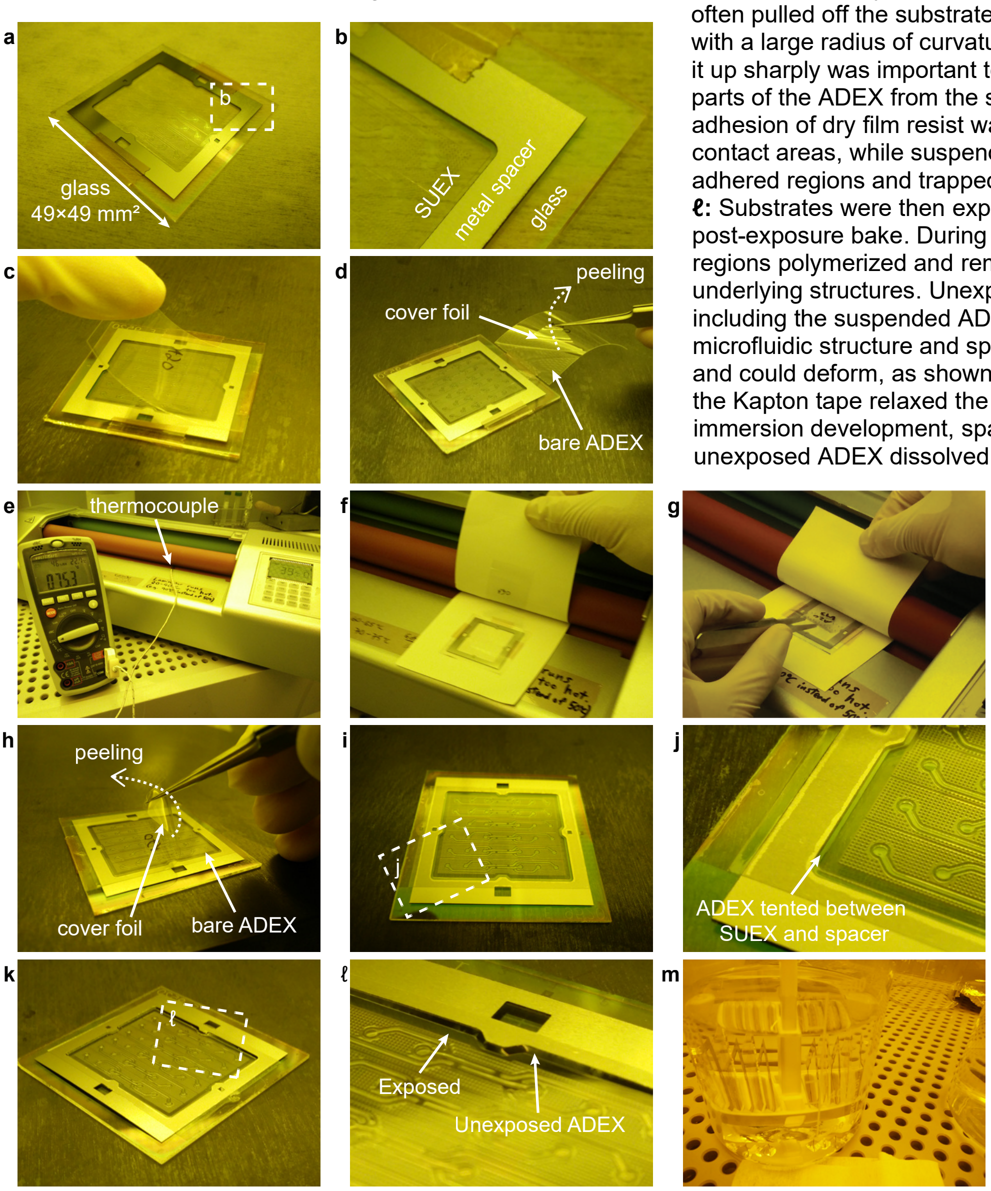

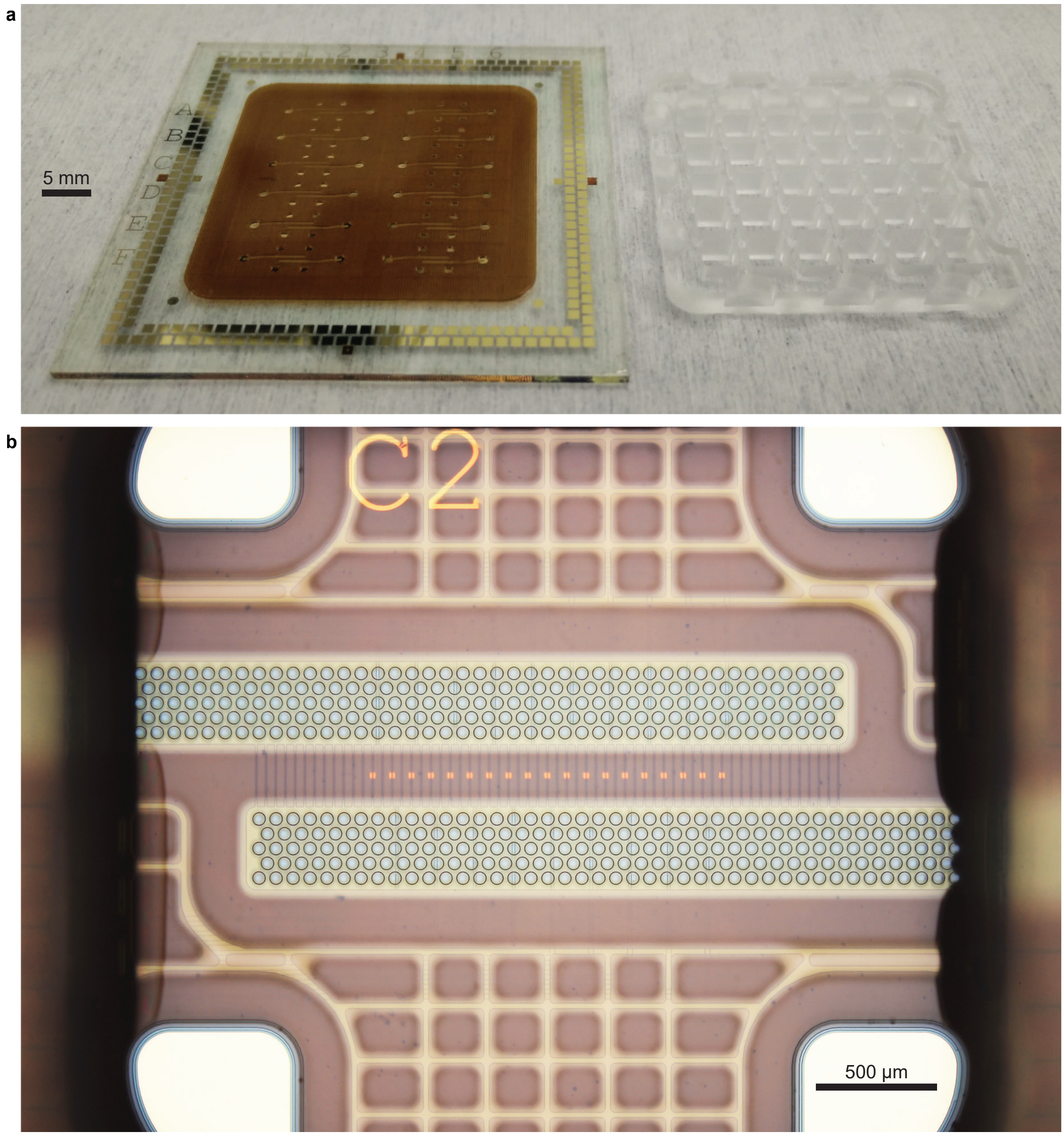


**Figure S3: Gluing of wells to microfluidics. a:** A microfluidic MEA after hard-baking (left) and a 6×6 well array (right) before gluing. For gluing, the well piece was manually aligned, and two-component epoxy was applied using a pipette at its edges. The glue wicked between the parallel surfaces by capillary action. **b:** Wicking stopped with a meniscus forming around each well, visible here on the left and right sides. The meniscus was pinned at the edge of features in the top ceiling layer, specifically the openings in the mesh ceiling (especially visible on the right side) and the openings above the reference electrode (four corners). The image is an uncropped version of **Figure 3g**.

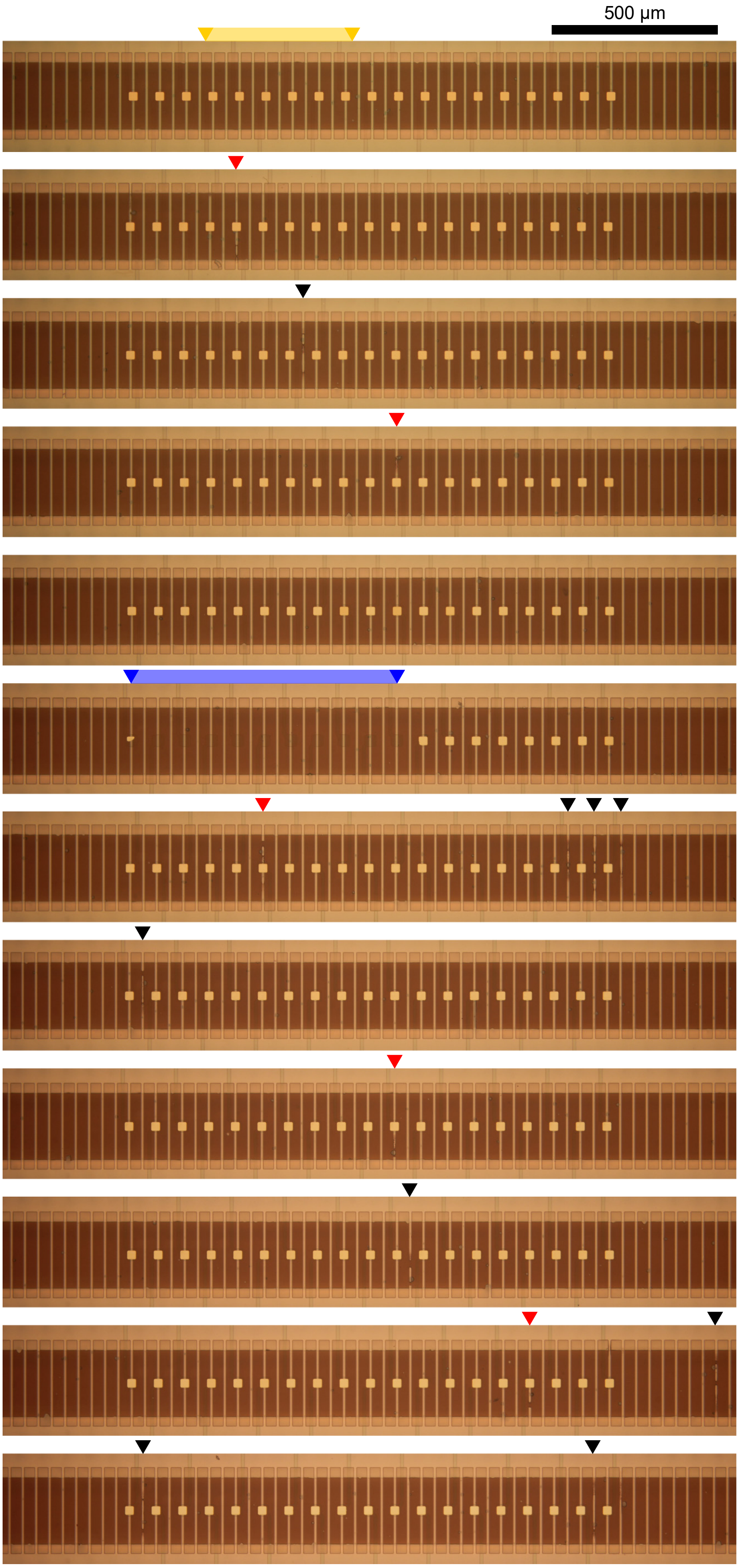


**Figure S4: Yield of tunnel fabrication.** All tunnels of a representative device are shown here. Approximately 2% of tunnels were observed to be blocked after hard baking. Out of 660 tunnels for 12 modules of a single device, 14 tunnels were blocked and are indicated by red arrows (five tunnels with electrodes) or black arrows (nine tunnels without electrodes). This device was representative, except for a defect during electrode manufacturing in the bottom left module (indicated in blue). Gold is missing on 11 electrodes, allowing an unobstructed view of the tunnels even above the structured electrodes. A representative section of the top module (indicated in yellow) is shown in **Figure 3e**.

A note on electrode defects: Typical yield of functional electrodes prior to microfluidic fabrication was 100 %. Here, we used “factory seconds” (MEAs with minor manufacturing defects). The device was specifically chosen due to the lithography defect which caused most gold electrode sites to be missing in the bottom left module. This allowed imaging from below of entire tunnels over the electrode region.

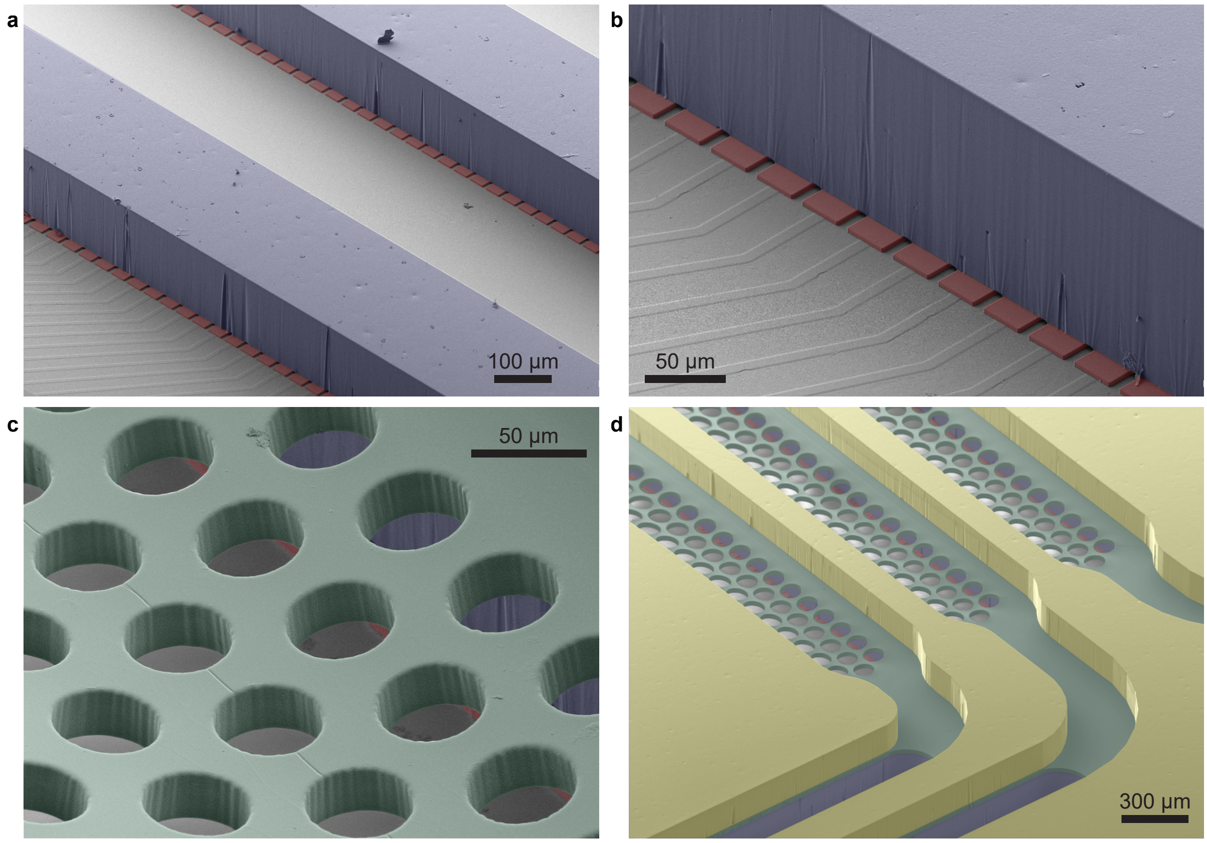


**Figure S5: False color scanning electron microscopy images of thick epoxy microfluidic structures. a, b:** SU-8 structures (red) define the 3 µm-tall tunnels and protrude from under the 150 µm-thick SUEX walls (blue) which define the compartments. The mesh ceiling was not yet applied. Topography of the insulated electrical paths is visible on the uncolored silicon nitride surface. **c:** A mesh ceiling (green, ADEX, 20 µm) covers the compartments, here with holes having a diameter of 50 µm as in the devices in this paper. **d:** In the design shown in this figure, a fourth layer (yellow, SUEX, 150 µm) defined channels above the mesh. Here, the mesh had a hole diameter of 100 µm. This figure shows a design having three adjacent compartments. The dimensions of the tunnel, compartment and mesh ceiling layers are the same here as in the two-compartment devices in this paper. The vertical walls of SUEX revealed vertical shadow-like defects, which may contribute to the tunnel blockages (**Figure S4**). Large areas were not segmented in this design (**d**).

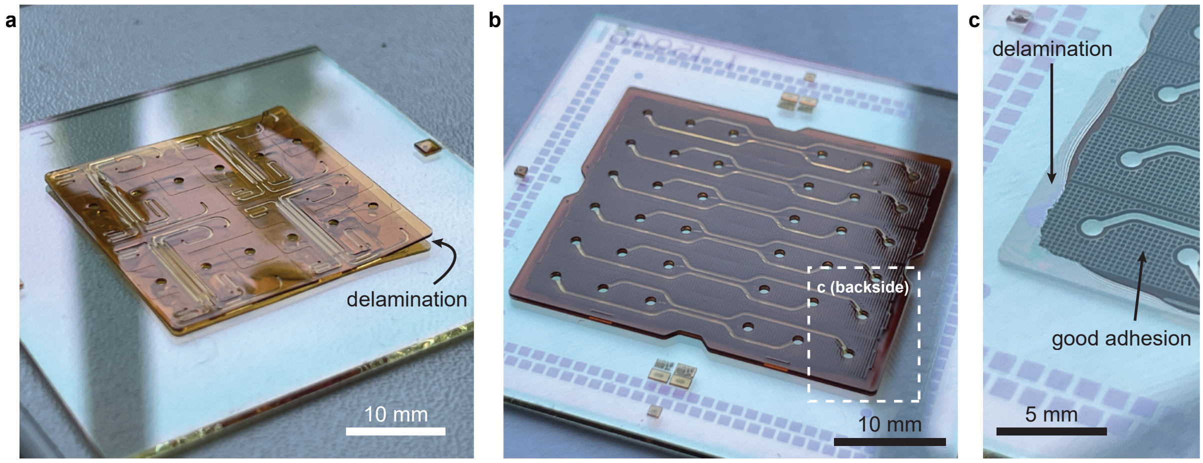


**Figure S6: Delamination of unoptimized epoxy microfluidics from glass.** Polymers experience much higher thermal expansion than glass. An early design (**a**) had a total epoxy thickness of 343 µm (nominal) and no segmentation of solid regions. Such structures massively delaminated after the final hard bake, even with slow ramping. Segmentation of solid regions significantly improved a later design (**b, c**) with the same 343 µm but delamination still occurred at ~1 mm-wide peripheral regions. Delamination is more easily visible from the backside (**c**), with clear contrast between dark well-adhered regions and bright delaminated regions with colorful interference patterns. The layers of these designs were (1) 3 µm SU-8, (2) 150 µm SUEX, (3) 20 µm ADEX, (4) 150 µm SUEX and (5) 20 µm ADEX. During processing, we observed that MEAs with layers 1–3 did not experience delamination, which supported the conception of the design described in this paper.

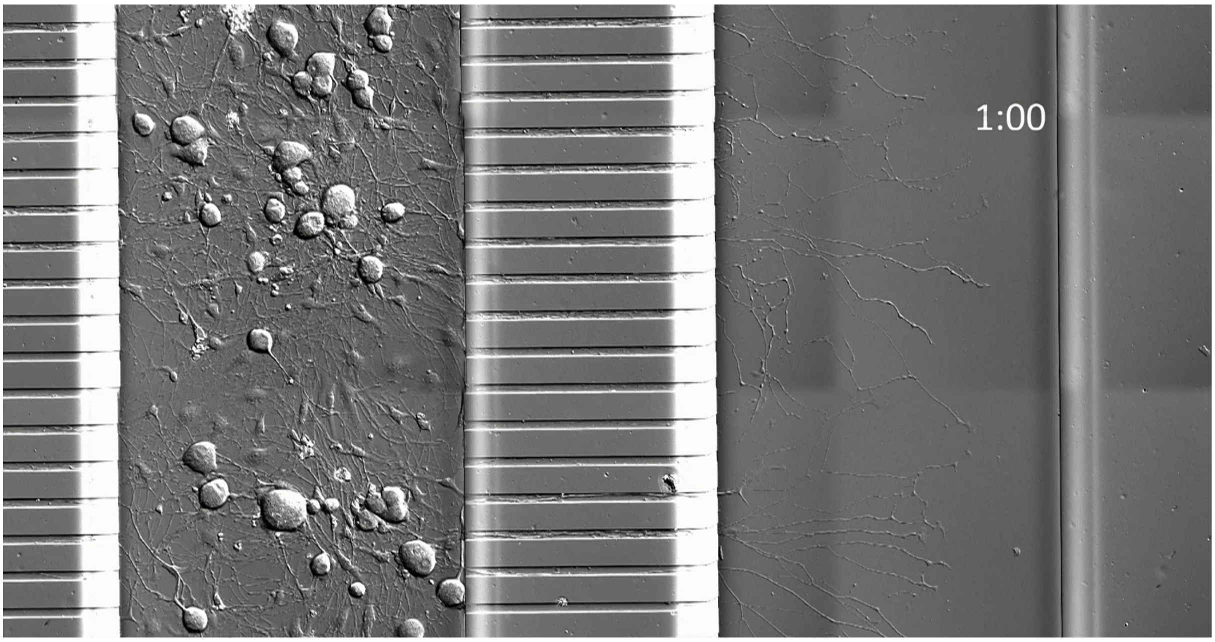


**Figure S7: Neurite outgrowth through tunnels (Supplementary Video 2).** Soma in the left compartment extended neurites in less than 24 h. For this time-lapse video over 48 h, dorsal root ganglion neurons were cultured in a PDMS microfluidic device and cultured adherent to the glass substrate. The use of adherent neurons rather than neurons suspended in hydrogel allowed live imaging by differential interference contrast microscopy. Time is indicated as days:hours.
